# Titers of antibodies the receptor-binding domain (RBD) of ancestral SARS-CoV-2 are predictive for levels of neutralizing antibodies to multiple variants

**DOI:** 10.1101/2022.03.26.484261

**Authors:** Trung The Tran, Eline Benno Vaage, Adi Mehta, Adity Chopra, Anette Kolderup, Aina Anthi, Marton König, Gro Nygaard, Andreas Lind, Fredrik Müller, Lise Sofie Nissen-Meyer, Per Magnus, Lill Trogstad, Siri Mjaaland, Arne Søraas, Karsten Midtvedt, Anders Åsberg, Andreas Barratt-Due, Asle W. Medhus, Marte Lie Høivk, Knut Lundin, Randi Fuglaas Karlsen, Reidun Dahle, Karin Danielsson, Kristine Stien Thomassen, Grete Birkeland Kro, Rebecca J. Cox, Fan Zhou, Nina Langeland, Pål Aukrust, Espen Melum, Tone Lise Åvitsland, Kristine Wiencke, Jan Cato Holter, Ludvig A. Munthe, Gunnveig Grødeland, Jan-Terje Andersen, John Torgils Vaage, Fridtjof Lund-Johansen

## Abstract

Diagnostic assays currently used to monitor the efficacy of COVID-19 vaccines measure levels of antibodies to the receptor-binding domain of ancestral SARS-CoV-2 (RBDwt). However, the predictive value for protection against new variants of concern (VOCs) has not been firmly established. Here, we used bead-based arrays and flow cytometry to measure binding of antibodies to spike proteins and receptor-binding domains (RBDs) from VOCs in 12,000 sera. Effects of sera on RBD-ACE2 interactions were measured as a proxy for neutralizing antibodies. The samples were obtained from healthy individuals or patients on immunosuppressive therapy who had received two to four doses of COVID-19 vaccines and from COVID-19 convalescents. The results show that anti-RBDwt titers correlate with the levels of binding- and neutralizing antibodies against the Alpha, Beta, Gamma, Delta, Epsilon and Omicron variants. The benefit of multiplexed analysis lies in the ability to measure a wide range of anti-RBD titers using a single dilution of serum for each assay. The reactivity patterns also yield an internal reference for neutralizing activity and binding antibody units per milliliter (BAU/ml). Results obtained with sera from vaccinated healthy individuals and patients confirmed and extended results from previous studies on time-dependent waning of antibody levels and effects of immunosuppressive agents. We conclude that anti-RBDwt titers correlate with levels of neutralizing antibodies against VOCs and propose that our method may be implemented to enhance the precision and throughput of immunomonitoring.

## INTRODUCTION

Clinical trials for COVID-19 vaccines showed that protection against symptomatic infection with ancestral SARS-CoV-2 (hereafter referred to as SARS-CoV-2wt) correlated with the levels of antibodies binding to the spike protein (spike-wt) and the receptor-binding domain (RBD-wt) ^1,2^. There is also evidence that neutralizing titers for SARS-CoV-2wt are predictive of protection against other variants including Delta ^3,4^. However, virus neutralization assays are poorly standardized and difficult to scale up ^3,5^. Thus, the neutralization titers corresponding to 90% vaccine efficacy against symptomatic COVID-19 varied by more than ten-fold in two clinical trials ^1,2^. Titers reported for binding antibodies (binding antibody units per milliliter, BAU/ml) after two doses of mRNA vaccines vary by seven-fold ^1,2,6-8^. There is therefore an unmet need for standardized high-throughput assays that can be used to monitor binding- and neutralizing antibodies against different SARS-CoV-2 variants at the individual level.

Most neutralizing antibodies interfere with the binding of the RBD to the human receptor ACE2 ^9-12^. Assays that measure inhibitory effects of serum on RBD-ACE2 interactions are therefore commonly used as a surrogate for virus neutralization assays ^8,10,13-17^. The optimal approach may be multiplexed measurement of ACE2-binding to RBDs from SARS-CoV-2 variants of concern (VOCs) ^8,15,18^. However, the methods are not standardized, and there is no consensus on the utility of RBD-ACE2 interaction assays in immunomonitoring.

Current diagnostic tests for humoral immunity against SARS-CoV-2 measure antibodies to RBD-wt. Results from a recent study suggest that this may be adequate since anti-RBD-wt titers were predictive of neutralizing activity of serum against the Delta and Omicron variants ^19^. Other studies, however, show that there is extensive person-to-person variation in ratios between binding- and neutralizing antibodies and in how mutations affect antibody binding and neutralization ^10,11,18^. It has also been suggested that the large increase in neutralization against the Omicron variant observed after a booster vaccine dose reflects an enhancement of antibody quality rather than in quantity ^20^. The implication of person-to-person variation in antibody quality would be that anti-RBDwt titers have limited predictive value at the individual level. However, to this end, studies on the qualitative variation in COVID-19 vaccine responses are on small cohorts, and little is known about effects of immunosuppressive therapy.

The aim of the present study was to determine if anti-RBDwt titers correlate with neutralizing antibodies as measured by RBD-ACE2 interaction assays. Arrays containing spike proteins and RBDs from SARS-CoV-2wt and the Alpha, Beta, Gamma, Delta, Epsilon, and Omicron BA.1 variants were incubated with 6693 sera (**Fig. 1**). In addition, more than 11.000 sera were analyzed with arrays containing proteins from all variants except Omicron. The arrays were labelled with fluorochrome-conjugated anti-human IgG to measure binding antibodies or with recombinant ACE2 to study effects of sera on RBD-ACE2 interactions (**Fig. 1**). In total, we analyzed 12,333 samples, and the cohort included COVID-19 convalescents, healthy individuals and patients on immunosuppressive therapy, who had received two to four doses of COVID-19 vaccines.

**Fig. 1.**
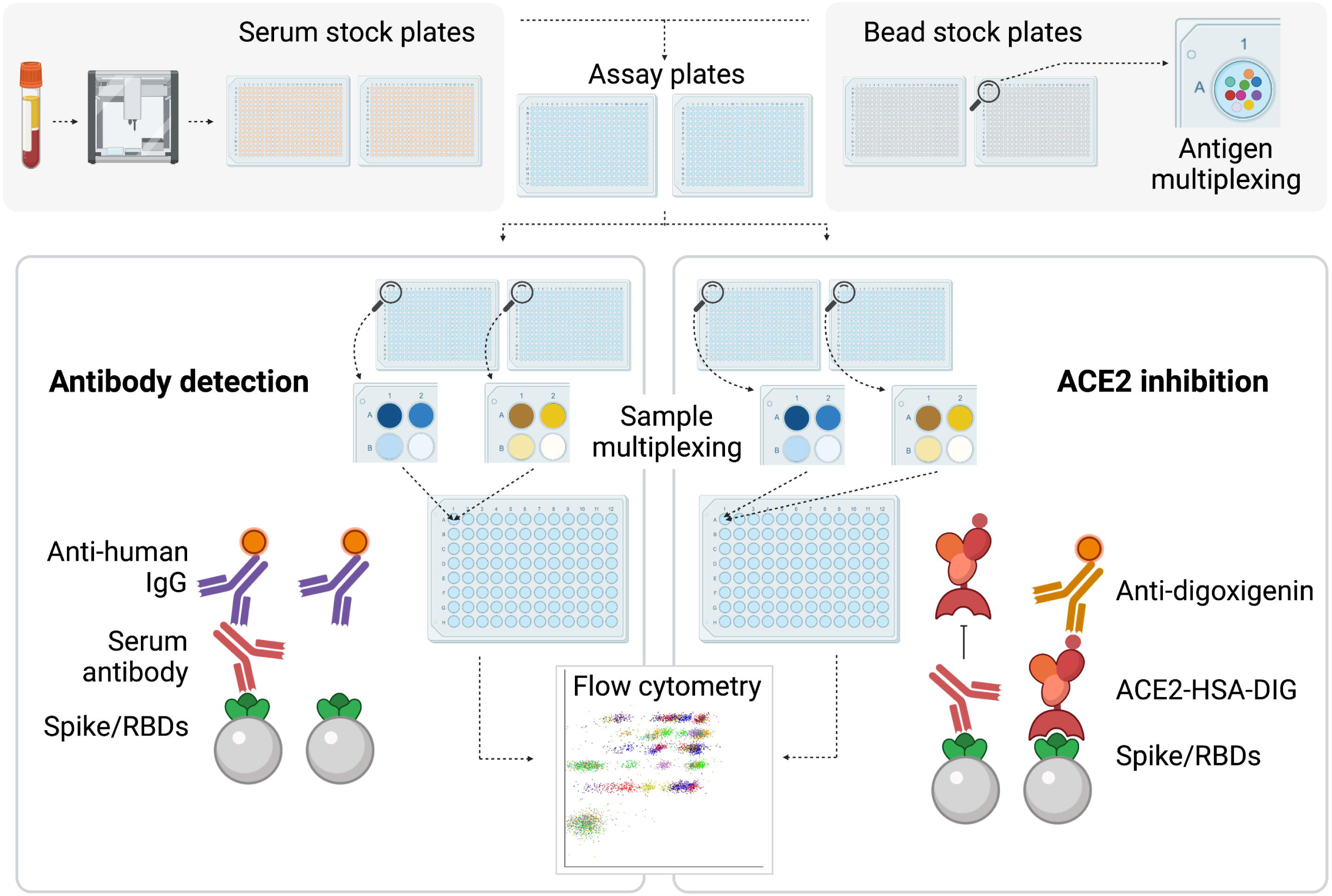
Multiplexed serology with high throughput. The figure shows an outline of the procedure. The starting point was the automated transfer of serum from blood collection tubes to two 384 well plates by use of a Tecan robot. Further processing was performed with liquid handling devices with 384 heads. Samples were serially diluted to 1:100 and 1:1000 in two pairs of 384 plates prefilled with assay buffer and bead-based arrays for measurement of RBD-ACE2 interactions and binding antibodies, respectively. Bead color codes corresponded to spike proteins and RBDs from SARS-CoV2 variants. After incubation with serum, the beads were washed and labelled with R-Phycoerythrin (PE)- conjugated anti-human IgG or successively with digoxigenin-conjugated ACE2 and anti-digoxigenin-PE. Eight additional barcodes were assigned to subarrays as addresses for positions A1-B2 in each of two 384 well plates. A robot with 96 heads was used to pool beads from one pair of 384 well plates into a single 96 deep well plate. Thus, each 96 well contained beads from eight samples for parallel analysis by flow cytometry.

## METHODS

### Serum samples

All participants gave informed consent before taking part in the study. Samples from Bergen were from studies approved by the Northern and Western Norway Regional Ethical Committees (approval numbers 218629, 118664). Samples from Oslo were obtained from the COVID-19 biobank at Oslo University Hospital. The biobank was approved by the Norwegian Regional Ethical Committee (reference number 135924). Sera from healthy volunteers were from participants in the Norwegian Coronavirus study, the Mother and Child study and the NorFlu study (approval numbers: 124170, 127798, 18403). Sera from kidney transplant recipients, patients with multiple sclerosis (MS), and inflammatory bowel disease (IBD) and other autoimmune diseases were obtained in context of ongoing studies on the immune responses to COVID-19 vaccination in the respective cohorts ^21-23^. All studies were approved by the Norwegian Regional Ethical Committee (reference numbers: 200631, 127798, 2021/8504, 135924, 204104).

### Expression and hapten-conjugation of recombinant ACE2

cDNA encoding a truncated human ACE2 fused to human albumin was sub-cloned into pFUSE2ss-CLIg-hk (InvivoGen). The vector was transiently transfected into Expi293F cells in suspension (Thermo Fisher Scientific) using the ExpiFectamine 293 transfection kit (Thermo Fisher Scientific) according to the manufacturer’s protocol. Cells were cultured for 7 days at 37°C with 80% humidity and 8% CO2 on an orbital shaker platform set to 125 rpm before the medium was collected. The secreted fusion protein was purified on a CaptureSelect™ human albumin affinity matrix (Life Technologies), and protein eluted by adding 20 mM Tris and 2.0 M MgCl2, pH 7.0 before up-concentration using Amicon® Ultra-15 50K Centrifugal Filter Units (Merck Millipore). Buffer exchange to PBS was performed before size exclusion chromatography (Äkta Avant, GE Healthcare) with a SuperdexTM 200 Increase 10/300 GL (Cytiva) prior to up-concentration using Amicon® Ultra-0.5 Centrifugal Filter Units (Merck Millipore). The protein eluted as a dimer.

### Expression and biotinylation of virus proteins

Except for Nucleocapsid (Prospec-Tany-TechnoGene, Israel) and RBD from Omicron (Sino Biologicals, China), all proteins were produced in-house. Plasmids encoding SARS-CoV2 RBD and full-length spike were obtained from Florian Krammer and Ian McLellan, respectively ^24,25^. The sequences were used as the basis for custom-made constructs encoding RBDs and Spike proteins from the Alpha, Beta, Gamma, Delta, and Epsilon variants (ordered from Genscript). cDNA encoding His-tagged Spike and RBD variants of SARS-CoV-2 were sub-cloned into pFUSE2ss-CLIg-hk (InvivoGen). The vectors were transfected as described for above the ACE2-albumin fusion protein and secreted His-tagged proteins were purified on HisTrap™ HP 1 mL columns (Cytiva), eluted with 250 mM imidazole diluted in PBS followed by up-concentration and buffer-exchange to PBS using Amicon® Ultra-15 10K Centrifugal Filter Units (Merck Millipore). Monomeric fractions were isolated by size exclusion chromatography as described above for the ACE2-albumin fusion protein. Purified recombinant viral proteins were solubilized in PBS and biotinylated chemically with sulfo-NHS-LC-biotin (sulfo-NHS-LC-biotin, Proteochem, USA) at a molar ratio of 1:1. Free biotin was removed with spin filters with a 10kDa cutoff (Merck, Millipore).

### Bead-based arrays with virus proteins

Bead-based arrays were produced using protocols described previously with some modifications ^26,27^. Amine-functionalized polymer beads (Bangs Laboratories, IN, USA) were suspended at 10% solids in PBS with 1% Tween 20 (PBT) in PCR-plates (Axygen). During all modification steps, the beads were agitated at 1800 rpm on an Eppendorf MixMate at 22°C. Each modification step was followed by five wash steps which include centrifugation of beads at 600 x g for 1 minute and resuspension of beads in PBT. Liquid handling was performed using CyBio SELMA robots with 96 heads (Jena Analytika, Germany). **Neutravidin coupling:** Beads were reacted successively with biotin-LC-NHS (sulfo-NHS-LC-biotin, 100µg/ml, Proteochem, USA) and neutravidin (Thermo Fisher, 100µg/ml). Each incubation lasted 30 min. **Fluorescent barcoding**: Beads were dyed successively with serially diluted Cy5-NHS (Lumiprobe), Bodipy-NHS (Lumiprobe), and Pacific Blue-NHS (Thermo Fisher) to generate a 108-plex (6 × 6 × 3). The starting concentrations were 100 ng-500 ng/ml, dilutions were 1:2.2, and the incubation time was 10-15 min. **Coupling of virus proteins**: Dyed Neutravidin-coupled beads were incubated for 30 min with biotinylated virus proteins solubilized in PBT (100 µg/ml). **Preparation and storage of bead-based arrays**: Beads with different color-codes and proteins were washed and then pooled in an assay buffer composed of PBT containing of 1% Bovine serum albumin (BSA), 0.1% sodium azide, 10 µg/ml D-Biotin, and 10 µg/ml Neutravidin. A production lot yields eight sub-arrays, each with 12 different barcodes and the same content of proteins. Ten color-codes corresponded to different virus proteins, while two were used as a reference for background binding of IgG to neutravidin beads. The eight sub-arrays have bar codes that can be discriminated by flow cytometry to allow sample multiplexing. They were distributed into positions A1, A2, B1, B2 in two 384 well plates prefilled with assay buffer. These served as stock plates and were kept at 4-8°C.

### Preparation of serum for analysis

Sample processing is outlined in **Fig. 1**. Serum (100 µl) was transferred from blood sampling tubes into 384 well serum stock plates using a Tecan Robot. A 384-head CyBio SELMA robot was used to transfer 10µl of serum into a 384 well plate prefilled with 90 µl serum dilution buffer. The buffer composition is the same as the assay buffer described above except that the neutravidin concentration is ten-fold higher (100 µg/ml) to neutralize neutravidin-reactive antibodies. The plates were kept overnight at 4-8°C before use. Serum remaining in the original 384 plates was stored at -20°C.

### Array-based measurement

The steps in the assay are illustrated in Fig. 1. A 384 head SELMA robot (Jena Analytica, Germany) was used to transfer beads (3 µl) from stock plates into two pairs of 384 well plates pre-filled with assay buffer. To one pair, we added 11 µl of diluted serum (1:10). The plate was subjected to mixing for 1 min on an Eppendorf MixMate before the beads were pelleted. Diluted serum (10 µl of 1:100 dilution) was next transferred to the second pair. The two plate pairs were agitated on an Eppendorf MixMate for one hour. At this point, the beads were pelleted and the supernatant was removed. For detection of IgG, the beads were washed three times in PBT and labelled for 30 min with R-Phycoerythrin-conjugated Goat-anti-Human IgG Fc (Jackson Immunoresearch, 30µl of a 1:600 dilution of stock in assay buffer). Beads used for ACE2-Spike interaction measurement were not washed. Digoxigenin-labelled recombinant ACE2 (30 µl, 300ng/ml) was added to the beads, and the plate was agitated for 40 min. The beads were washed twice in PBT and labelled with monoclonal anti-Digoxin (Jackson Immunoresearch, 1µg/ml) conjugated in-house to R-Phycoerythrin for 30 min at constant agitation. After labeling with secondary antibodies, the beads were washed twice. The bead-based arrays in positions A1, A2, B1, B2 in each 384 well plate have different barcodes. Thus, the contents of the two 384 well plates were pooled into a single 96 well deep well plate prior to flow cytometric analysis. The beads were next analyzed with an Attune Next Flow cytometer equipped with four lasers (violet, blue, yellow, and red) and a harvesting unit for microwell plates). Instrument usage averaged 60 minutes per 96 well plate.

### Data analysis and visualization

Raw flow cytometry data were analyzed using WinList 3D version 10. The median R-Phycoerythrin fluorescence intensity (MFI) for each bead subset was exported to a spreadsheet. Further analysis was performed in Microsoft Excel. The MFI values measured for binding of anti-human IgG and ACE2/anti-digoxigenin to beads with viral proteins were divided by those of beads with neutravidin only (hereafter referred to as relative MFI, rMFI). To visualize results as colored dot plots, we exported data from Excel to WinList 3D. In all dot plots in Fig.1-9 and supplementary figures, each dot corresponds to a single sample. Colors correspond to subgroups of samples as indicated in the text and figure legends.

### Calibration of signals to Binding Antibody Units per milliliter (BAU/ml)

Serum was obtained from a healthy individual two weeks after receiving an mRNA vaccine booster dose. The serum was serially diluted and analyzed using the Roche Elecsys anti-SARS-CoV-2 S assay. The end titer was 53.000 BAU/ml. The serum stock was diluted serially two and three-fold to generate a standard series in the range of 3-50.000 BAU/ml, and the standard series was added to separate wells in all 384 plates analyzed. Numerical values for rMFI measured for anti-RBD-wt and RBD-ACE2 interactions were subjected to regression in Excel (**supplementary methods**). The best curve-fit was obtained with the power function (i.e. calculation based on log-transformed data). The signals measured with study samples were used as input in regression functions to convert rMFI to BAU/ml. See also **supplementary methods**.

### Statistics

Pearson correlations for log-transformed data were determined using Excel.

### Virus neutralization assay

#### Laboratory 1 Oslo

The virus neutralization assay has been described previously ^5^. Vero E6 cells were plated out into 96-well cell culture plates at 1×104 cells/well. The next day, 100xTCID of SARS-CoV-2 virus were mixed with 2-fold titrations of sera assayed in quadruplicates. Following one hour incubation at 37°C in a 5% CO_2_ humidified atmosphere, the mixture was added to the plated cells. Plates were next incubated for 50 hours at 37°C in a 5% CO_2_ humidified atmosphere. Next, monolayers were washed with PBS and fixed in cold 80 % acetone for 20 min.

The SARS-CoV-2 virus Human 2019-nCoV strain 2019-nCoV/Italy-InMI1 (008V-03893) from the European Virus Archive (EVA) was detected in Vero E6 cell cultures by ELISA using a mAb against SARS-CoV-2 nucleocapsid (40143-R004, SinoBiological) and HRP-conjugated goat anti-rabbit IgG-Fc mAb (SSA003, SinoBiological). Plates were developed using TMB substrate buffer (sc-286967, Santa Cruz), and read with the Tecan reader.

#### Laboratory 2: Bergen

The microneutralization (MN) assay was performed in a certified Biosafety Level 3 Laboratory in Norway^17-19^ against a clinically isolated virus: SARS-CoV-2/Human/NOR/Bergen1/2020 (GISAID accession ID EPI_ISL_541970) ^28^. Briefly, serum samples were heat-inactivated at 56°C for 60 min, analysed in serial dilutions (duplicated, starting from 1:20), and mixed with 100 TCID_50_viruses in 96-well plates and incubated for 1 hour at 37° C. Mixtures were transferred to 96-well plates seeded with Vero cells. The plates were incubated at 37°C for 24 hours. Cells were fixed and permeabilized with methanol and 0.6% H2O2 and incubated with rabbit monoclonal IgG against SARS-CoV2 NP (Sino Biological). Cells were further incubated with biotinylated goat anti-rabbit IgG (H+L) and HRP-streptavidin (Southern Biotech). The reactions were developed with o-Phenylenediamine di-hydrochloridec (OPD) (Sigma-aldrich). The MN titer was determined as the reciprocal of the serum dilution giving 50% inhibition of virus infectivity. Negative titers (<20) were assigned a value of 5 for calculation purposes.

##### Enzyme-linked immunosorbent assay (ELISA)

SARS-CoV-2 specific antibodies were detected using an ELISA as previously described but with some modifications ^24,28^. Sera were screened for IgG antibodies against the receptor-binding domain (RBD, Wuhan) of the SARS-CoV-2 Spike protein at a 1:100 dilution and the samples were run in duplicates. The diluted sera were incubated for 2 hours at room temperature in 96 well plates (Maxisorp, Nunc, Roskilde, Denmark) coated with 100 ng/well of the RBD antigen. Bound IgG antibodies were detected with a horseradish peroxidase (HRP)-labeled secondary antibody (cat. no.: 2040-05, SouthernBiotech, Birmingham, AL, USA) and the addition of the chromogenic substrate 3,3′,5,5′-tetramethylbenzidine (TMB; BD Biosciences, San Jose, CA, USA). Optical density (OD) was measured at 450/620 nm using the Synergy H1 Hybrid Multi-Mode Reader with the Gen5 2.00 (version 2.00.18) software (BioTek Instruments Inc., Winooski, VT, USA).

RBD positive sera were run in an additional ELISA, where the ELISA plates were coated with SARS CoV-2 Spike protein (Wuhan, 100ng/well). Sera were serially diluted in duplicate in a 5-fold manner, starting from a 1:100 dilution, and the ELISA plates were incubated with diluted serum for 2 hours at room temperature. Bound IgG antibodies were detected and measured as described for the RBD screening ELISA.

Positive controls were serum from a hospitalized COVID-19 patient and CR3022 ^29^, whereas pooled pre-pandemic sera (n=128) were used as a negative control ^28^. The endpoint titers were calculated for each sample. Samples with no detectable antibodies were assigned a titer of 50 for calculation purposes.

## RESULTS

### Antibodies in sera from COVID-19 convalescents and vaccinated individuals have broad and uniform coverage of RBDs from SARS-CoV-2 variants

We used bead-based arrays and flow cytometry to measure IgG antibodies to Nucleocapsid (wt) and RBDs from SARS-CoV-2 variants in 5145 sera diluted 1:1000 (**Fig. 2**). The samples were obtained in 2020 from COVID-19 convalescents (n=318, red dots), or in 2021 or 2022 from vaccinees (green dots). The post-vaccine samples were from healthy individuals (n=1060), immunocompetent individuals who had tested positive for the Delta variant (n=43) and patients on immunosuppressive therapy (3703). We also included 23 samples collected from a cohort of double-vaccinated immunocompetent individuals with Omicron infection (blue dots).

**Fig. 2.**
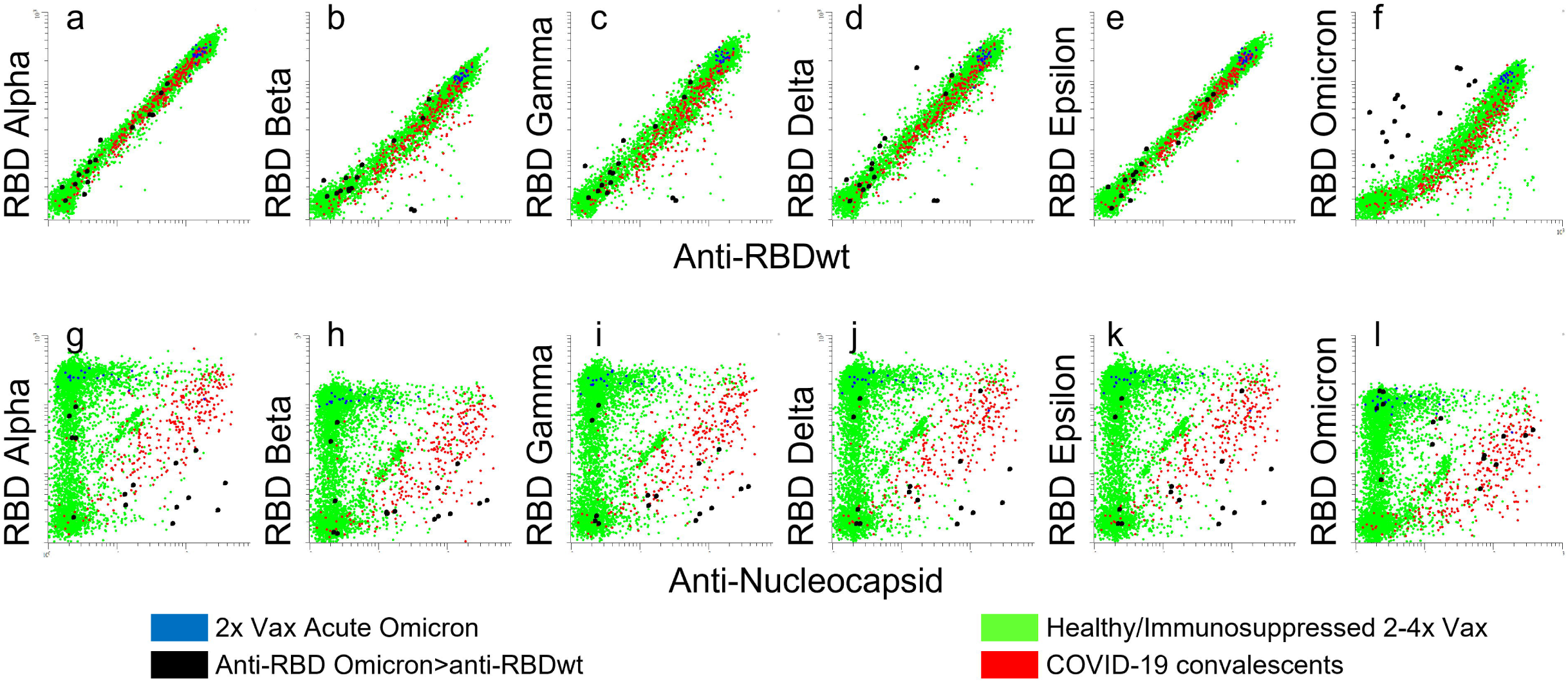
Correlation between levels of antibodies to RBDs from SARS-CoV-2 variants. The dot plots show accumulated data from array-based measurement of 6693 samples. Each dot in the scatter plots correspond to a different sample. Bead-based arrays were incubated with 6693 sera diluted 1:1000 for 1h prior to labeling with R-Phycoerythrin-conjugated anti-Human IgG. **a-f:** The x-axes show median fluorescence intensity (MFI) of anti-human IgG binding to beads coupled with RBD-wt divided by MFI measured for beads with no virus protein (relative MFI, rMFI). The y-axes show anti-human IgG rMFI measured for beads coupled with RBDs from indicated variants. **g-l:** The y-axes are the same as in dot plots a-f, while the x-axis shows rMFI for antibodies binding to beads coupled with nucleocapsid from SARS-CoV-2wt. Green dots: post-vaccine sera, red dots: sera obtained in 2020 from SARS-CoV-2 convalescents. Blue dots: sera obtained 10-20 days after symptom debut of Omicron infection in double-vaccinated healthy individuals.

The overall correlation (Pearson correlation coefficients) with levels of antibodies to RBD-wt was 0.91 for RBD from Omicron and 0.94 or higher for antibodies to RBDs from all other variants tested (**Fig. 2a-f**). In samples obtained during 2020 from COVID-19 convalescents, the correlation between anti-RBD-wt and anti-Nucleocapsid was 0.56 (**Fig. 2**, red dots). Reactivity with Omicron RBD was lower in convalescent sera, and the correlation with anti-RBD-wt was 0.79 (**Fig. 2f**, red dots, **supplementary Fig. 1**).

We identified 14 samples with stronger binding of IgG to RBD from Omicron than to RBD-wt (**Fig. 2, black dots**). All were from 2022, and 9 contained antibodies to nucleocapsid (**Fig. 2g-l**). The samples are therefore likely to be from SARS-CoV-2/vaccine naive individuals infected with Omicron ^30^. Overall, signals measured for binding of antibodies to RBD from Omicron were weaker than those measured for RBDs from other variants (**Fig. 2f, supplementary Fig. 1**). This was also observed in sera obtained from double-vaccinated individuals 12-20 days symptom debut of Omicron infection (**Fig. 2, blue dots**).

To extend the dynamic range of antibody detection, we analyzed 438 sera at dilutions of 1:10,000 and 1:100,000. The correlations were similar to those observed after measurement at 1:1000 dilution (**supplementary Fig. 2**). Collectively, these results show that most individuals generate antibodies with broad and similar coverage of RBDs from SARS-CoV-2 variants.

### The relative content of neutralizing antibodies against different SARS-CoV-2 variants is similar in COVID-19 convalescents and vaccines

RBD-ACE2 interactions were measured as a surrogate for neutralizing antibodies. The sera were diluted 1:100, incubated with bead-based arrays and labelled with recombinant ACE2 (**Fig.1**). We identified four groups of anti-RBD-wt-positive sera (**Fig. 3a-b**). I) content of antibodies with minimal inhibition of ACE2-RBD interactions (**red**,17.6%), II) inhibition of ACE2-binding to RBD-wt (**green**, 33%), III) complete inhibition of ACE2-binding to RBD-wt and/or partial inhibition of binding to RBD-Beta (**orange**, 27%), and IV) complete inhibition of ACE2 binding to RBD-Beta (**blue**, 22%).

**Fig. 3.**
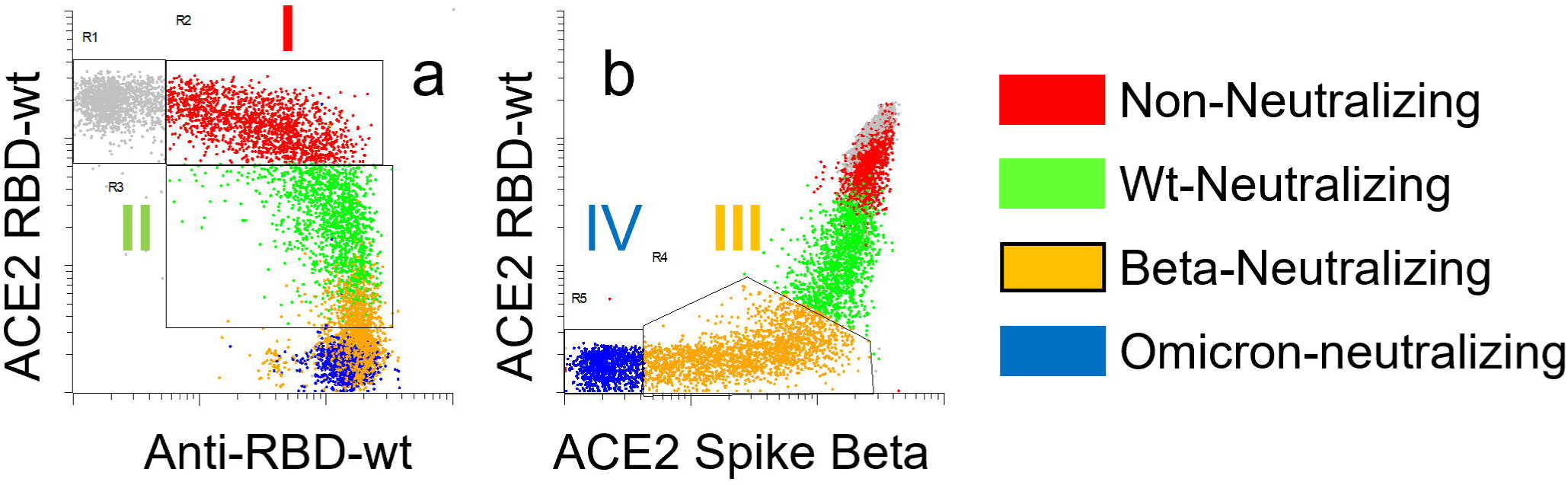
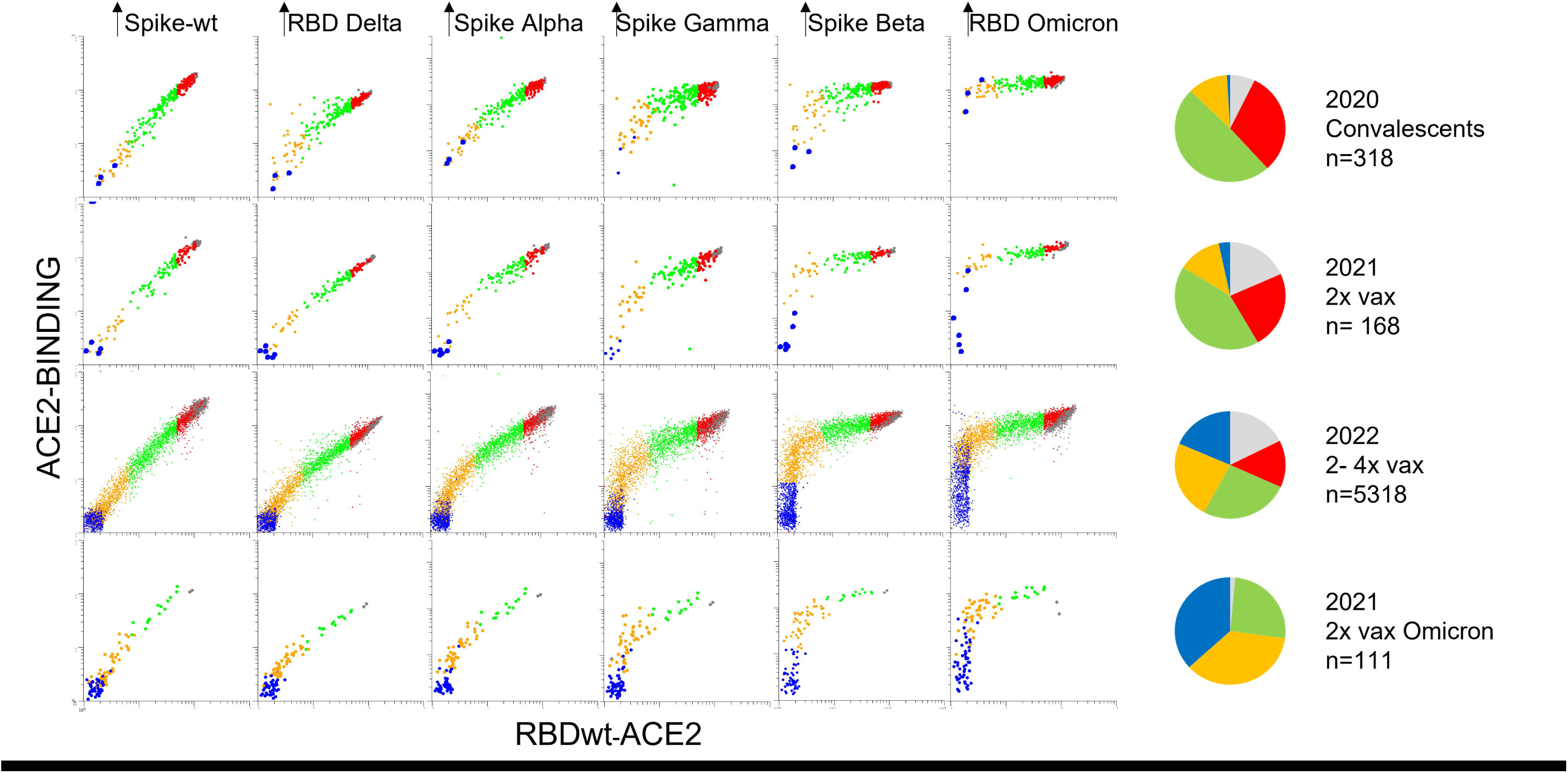
Effects of serum on ACE2-binding to RBDs and spike proteins from SARS-CoV-2 variants. The dot plots show accumulated data from array-based measurement of 6693 samples. Each dot in the scatter plots correspond to a different sample. **a-b:** Binding of serum IgG or recombinant ACE2 to beads coupled with RBD-wt or RBD from the Beta variant as indicated. The beads were incubated with serum diluted 1:1000 or 1:100 prior to labeling with anti-human IgG or recombinant ACE2, respectively. **c-h**. Binding of ACE2 to beads coupled with RBDs and spike proteins from indicated SARS-CoV-2 variant. The colors correspond to groups within the regions shown in panels **a** and **b**. Grey dots anti-RBD-wt negative sera. Red dots: anti-RBD-positive sera with no or minimal inhibition of ACE2-binding to RBD-wt. Green dots: sera with selective and partial inhibition of ACE2-binding to RBD-wt. Orange dots: sera with complete inhibition of ACE2-binding to RBD-wt and partial inhibition of binding to RBD-Beta. Blue dots. Complete inhibition of ACE2-binding to RBD-wt and RBD-beta.

The inhibitory effects of serum on ACE2-binding to RBDs from SARS-CoV-2 variants followed a stringent and uniform pattern in sera from all cohorts (**Fig. 3c**). Effects on ACE2-binding to RBD-wt and RBD-Delta were similar, while the resistance against serum inhibition for other variants increased from Alpha, Gamma, Beta to Omicron (**Fig. 3c**). The differences between the cohorts were in the distribution of samples in each of the groups identified in **Fig. 3a-b**. Thus, there was an increase in the frequency of sera with neutralizing antibodies against all variants starting from samples obtained in 2020 from COVID-19 convalescents to those obtained from double vaccinated individuals with acute Omicron infection (**Fig. 3c**). Collectively, these results show that there is little interindividual variation in the relative content of neutralizing antibodies to different SARS-CoV-2 variants.

From here on, the groups are referred to as non-neutralizing (red), wt-neutralizing (green), Beta-neutralizing (orange) and Omicron-neutralizing (blue).

### Anti-RBD-wt titers are predictive for the levels of neutralizing antibodies to all SARS-CoV-2 variants

We aligned results from RBD-ACE2 interaction assays with those obtained by measuring anti-RBDwt in sera diluted 1:100,000 (**Fig. 4**). The plots in **Fig. 4c-g** show that there was an inverse correlation between ACE2-binding and anti-RBDwt titers. However, there were variant-specific thresholds for the inhibitory effect. Thus, anti-RBDwt signals showing linearity with ACE2-binding to RBD from Omicron (y-axis) were right-shifted by approximately one log compared those with co-linearity with ACE2 binding to RBDwt (**Fig. 4c-g**).

**Fig. 4.**
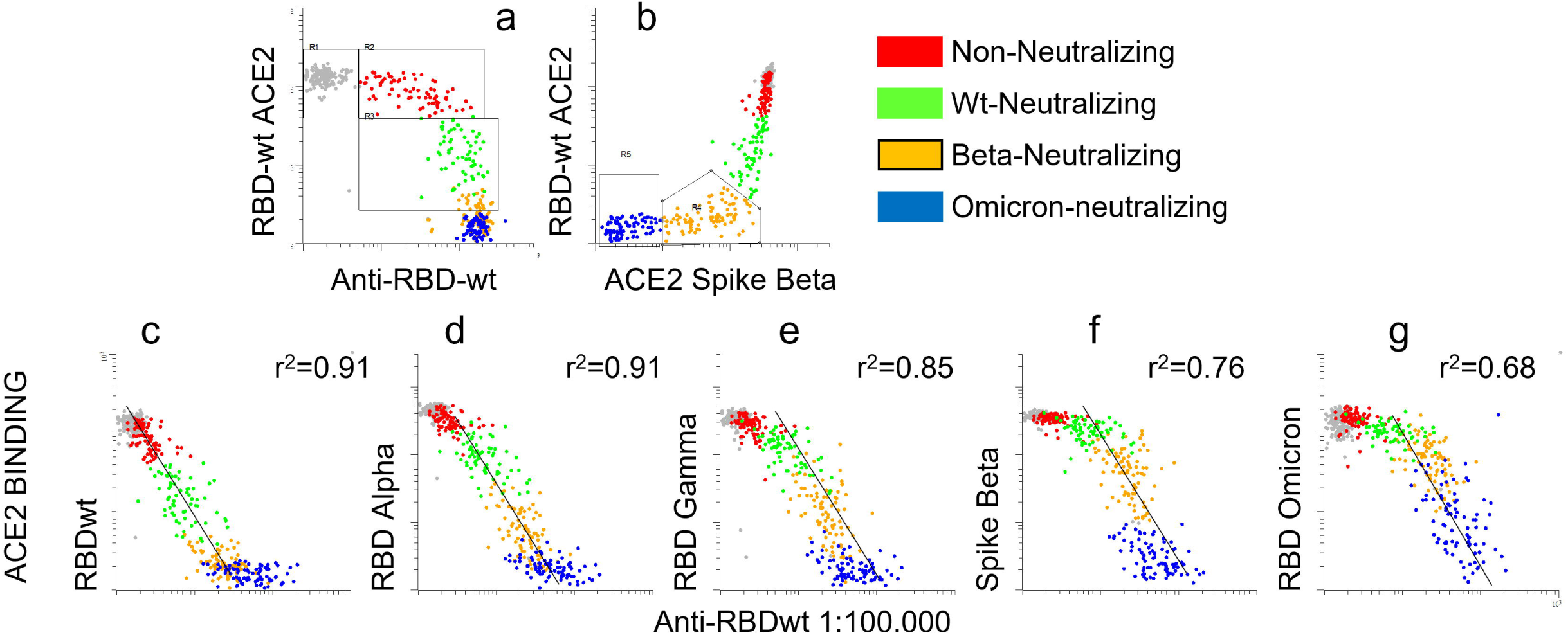
Inhibitory effects of RBD-ACE2 interactions correlate with anti-RBD-wt titers. **a-b:** see legend to Fig.3a-b. Each dot in the scatter plots correspond to a different sample. **c-g:** The x-axes show binding of human IgG to RBDwt after incubation with serum diluted 1:100,000. The y-axes show binding of ACE2 to beads coupled with RBDs and spike proteins from indicated SARS-CoV-2 variant after incubation with serum diluted 1:100. The dot plots show accumulated data from array-based analysis of 550 sera. Each dot corresponds to a different sample. Pearson correlation coefficients were determined for log-10 transformed data.

Sera obtained in 2020 from COVID-19 convalescents and a subset of those obtained in 2021 from double-vaccinated individuals had been measured with an in-house anti-spike ELISA earlier. There was an inverse relationship between anti-spike titers and RBD-ACE2 interactions (**Fig. 5**). Inhibition of ACE2-binding to Omicron RBD corresponded to titers higher than 10^5^ (**Fig. 5l, blue dots**). These were from individuals who had recovered from COVID-19 in 2020 and later received one vaccine dose in 2021. Collectively, the results in **Fig.4-5** show that anti-RBD-wt titers are predictive for the content of neutralizing antibodies to SARS-CoV2 variants.

**Fig. 5.**
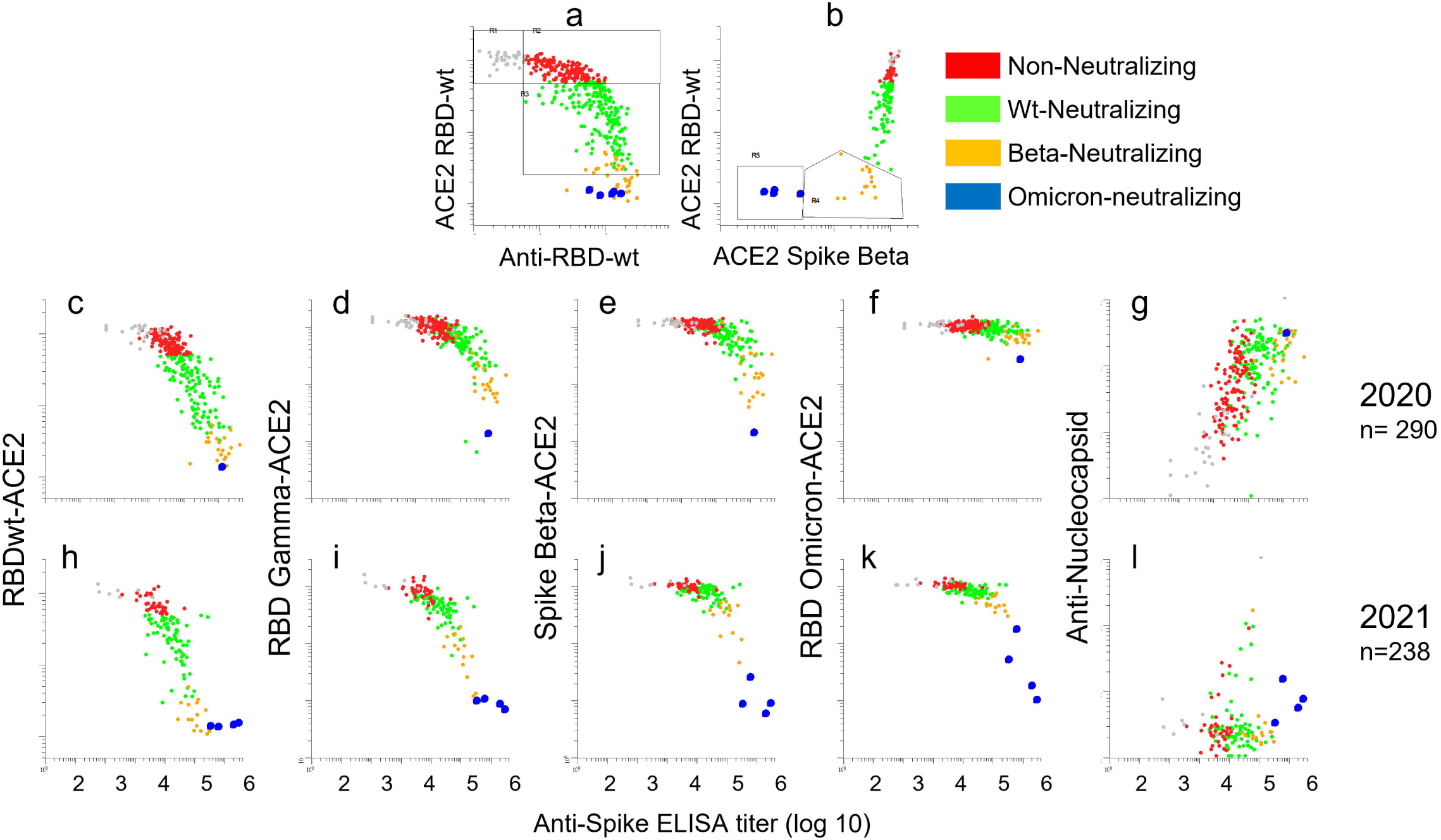
Inhibitory effects of RBD-ACE2 interactions correlate with anti-spike wt titers as measured by ELISA. A total of 428 sera that had been measured with an in-house ELISA for anti-spike wt were analyzed with Multi-IgG-ACE2-RBD. Each dot in the scatter plots correspond to a different sample. **a-b:** see legend to Fig.3a-b. **c-l:** the y-axes show binding of ACE2 to beads coupled with RBDs and spike proteins from indicated SARS-CoV-2 variant after incubation with serum diluted 1:100. The x-axes show anti-spike wt titers as measured by ELISA. 2020: Sera obtained in 2020 from COVID-19 convalescents. 2021: sera obtained in 2021 from COVID-19 convalescents and double vaccinated healthy individuals.

### Multi-IgG-ACE2-RBD yields an internal reference for binding antibody units per milliter (BAU/ml)

The assay plates used to generate the results shown in **Fig. 2-5** contained a standard series that was calibrated to BAU/ml using the Roche Elecsys anti-SARS-CoV-2 S assay (see methods). Results obtained with the standard series in 26 consecutive 384 well plates are highlighted as black dots in **Fig. 6 a-b** and colored according to the parent groups in **Fig. 6c-f**. Signal values measured for standards were subjected to regression in Excel to generate formulas for conversion of signals from test samples to BAU/ml (**Fig. 6g-j, supplementary methods)**.

**Fig. 6.**
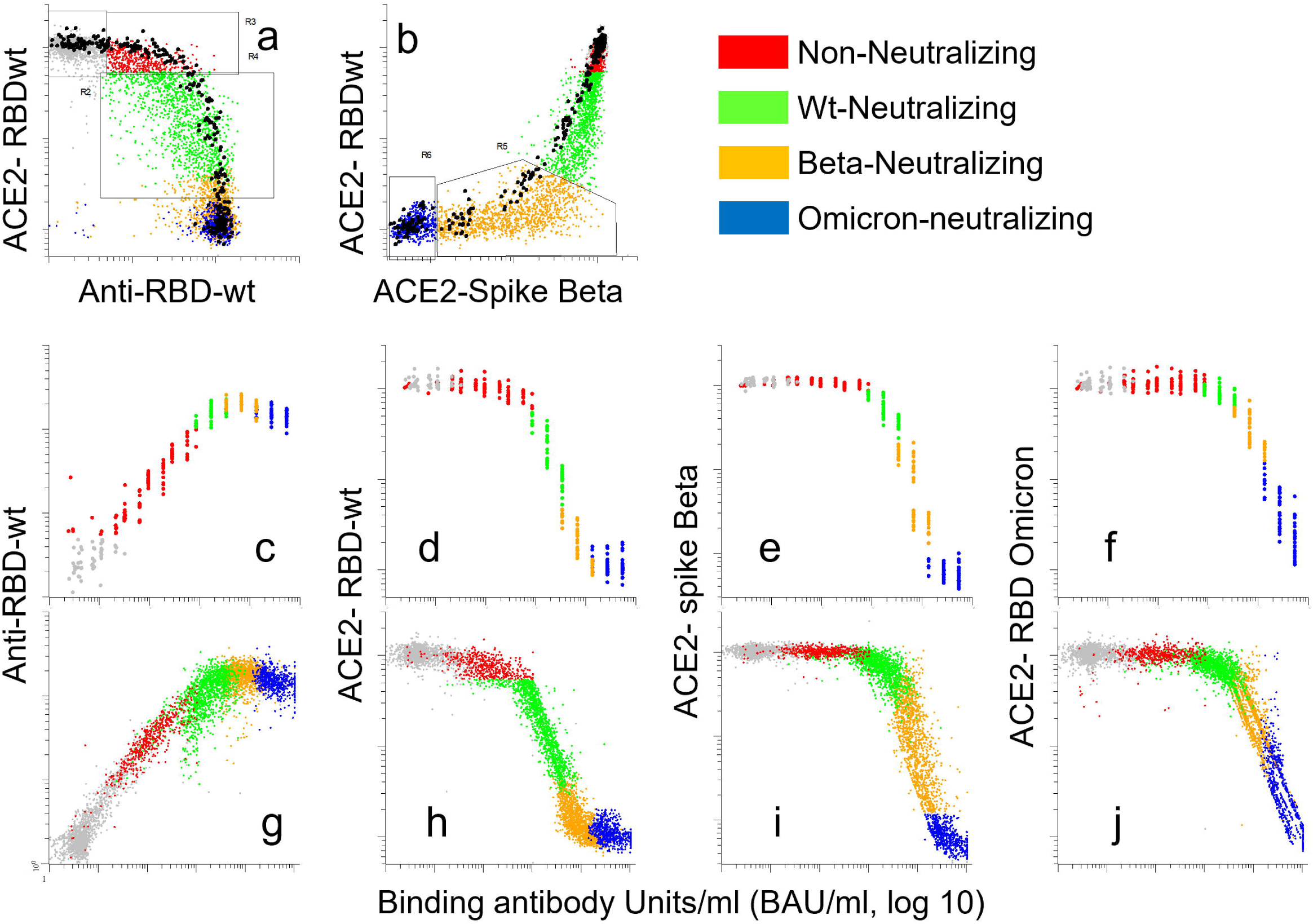
Groups identified by multi-IgG-ACE2-RBD correspond to defined ranges of binding antibody units per milliliter (BAU/ml). A total of 6693 sera were analyzed with multi-IgG-ACE2-RBD together with a standard series with pre-defined BAU/ml. Each dot in the scatter plots correspond to a different sample. **a-b:** see in the legend to Fig.3a-b, except that the black dots correspond to a standard series prepared by serial dilution of a serum with an anti-RBD-wt titer of 53.000 BAU/ml. **c-f:** Results obtained with the standard series in 26 consecutive 384 well plates. The x-axes show BAU/ml calculated from serial dilutions of the standard series. The y-axes show binding of IgG to beads coupled with RBD-wt (**c-g**), or ACE2 binding to beads coupled with RBDs or spike proteins as indicated. **g-j**: Results obtained with the standard series were used as input in Excel regression functions to generate formulas for conversion of signals measured for the 6693 sera into BAU/ml (supplementary methods).

The results in **Fig. 6** show that non-neutralizing sera contained 30-500 BAU/ml (**Fig. 6 c-f, red dots**), wt-neutralizing corresponded to 500-3000 BAU/ml (**Fig. 6 c-f, green dots**), beta-neutralizing to 3000-11.000 BAU/ml while Omicron-neutralizing sera contained more than 11.000 BAU/ml (**Fig. 6d-f**). Reactivity patterns obtained by Multi-IgG-ACE2-RBD therefore yield an internal reference for anti-RBD-wt titers.

### Multi-IgG-ACE2-RBD yields an internal reference for neutralizing activity of serum against SARS-CoV-2wt

Multi-IgG-ACE2-RBD was used to analyze sera that had been tested for neutralizing activity against SARS-CoV-2wt in two different laboratories (lab1: n=364, lab2: n=138). The correlation coefficients between ACE2-binding to RBD-wt and neutralization titers were -0.92 and -0.89, respectively (**Fig. 7**). The cutoff for neutralizing activity in laboratory 1 was a titer of 20, which corresponded to approximately 500 BAU/ml (**Fig. 7e**), and 77% of samples with titers higher than 20 in laboratory 2 also contained at least 500 BAU/ml (**Fig. 7j**). The titers for Omicron-neutralizing sera were at the upper limit of the assay (**Fig. 7d**).

**Fig. 7.**
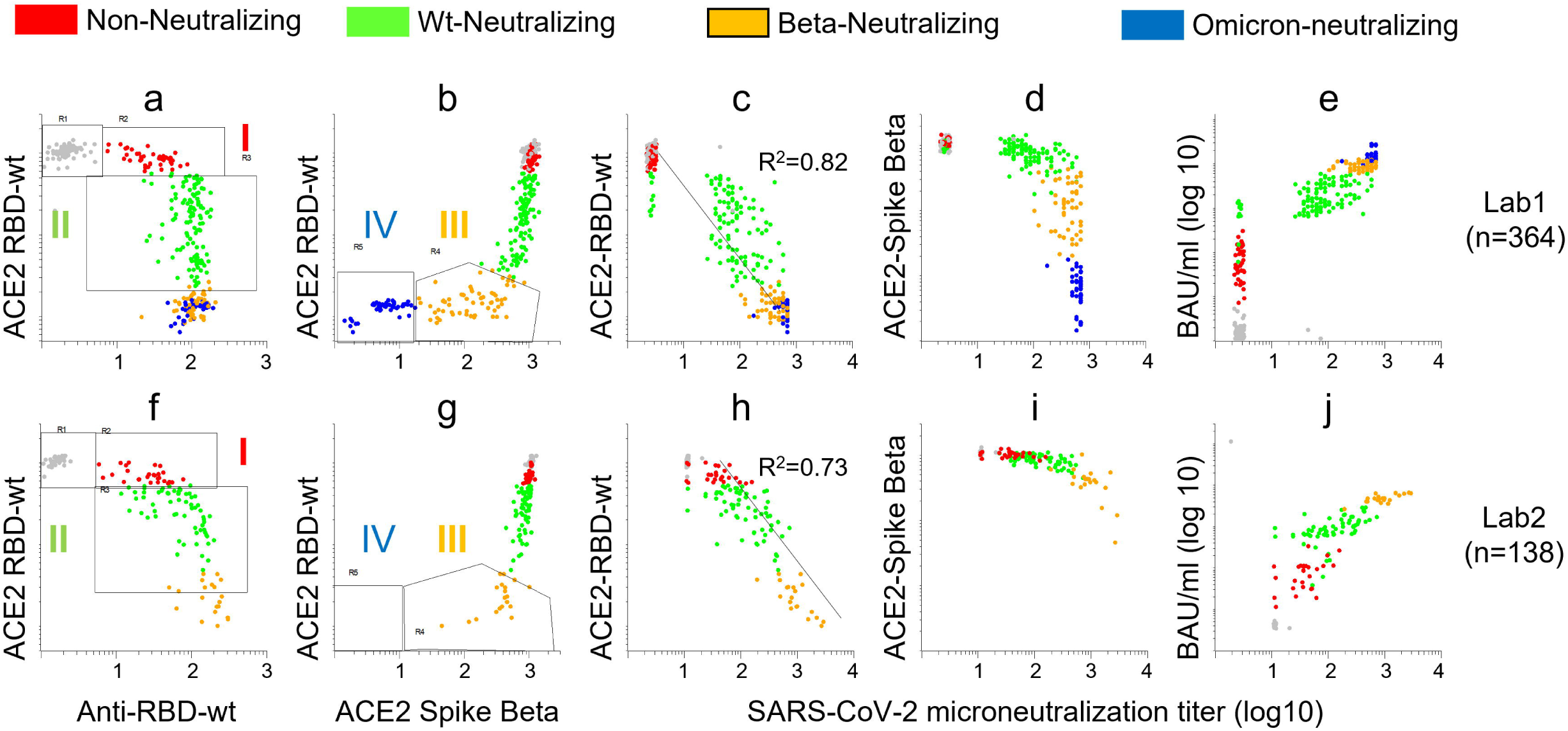
Groups identified by multi-IgG-ACE2-RBD correspond to defined ranges of neutralizing activity against SARS-CoV-2wt. A total of 502 sera that had been tested for neutralizing activity against SARS-CoV-2 were analyzed with Multi-IgG-ACE2-RBD. Each dot in the scatter plots corresponds to a different sample. **a-b and f-g:** see legend to Fig.3a-b, **c-e**: the x-axis shows virus neutralization titers against SARS-CoV-2wt measured for 364 sera in Oslo. The y-axes show ACE2-binding to RBD-wt (c) or spike-Beta (d) while panel **e** shows RBD titers as BAU/ml. h-j: the x-axis shows virus neutralization titers against SARS-CoV-2wt measured for 138 sera in Bergen.

### Results from Multi-IgG-ACE2-RBD analysis recapitulate published knowledge about time-dependent waning of antibodies and effects of immunosuppressive therapy

During 2021, we used Multi-IgG-ACE2-RBD to monitor the effects of COVID-19 vaccination of healthy individuals and patients on immunosuppressive therapy. At that time, we did not have access to the Omicron RBD. The upper dynamic range of the assay was therefore approximately 20.000 BAU/ml.

Among sera obtained from healthy individuals 10-50 days after the second vaccine dose, 98% were classified as neutralizing (i.e. >500 BAU/ml), and 70% as Beta-neutralizing (**Fig. 8c, green, orange, and blue dots, respectively**, see also pie charts in **supplementary Fig. 3**). The median titer was 6425 BAU/ml. Our assay was calibrated against the Roche Elecsys anti-SARS-CoV-2 S assay, and the titers measured here were comparable to those reported for the reference assay earlier ^7^. The time-dependent waning of antibody levels was also in line with results reported earlier ^31^. Thus, four months after vaccination, the median titer was reduced by approximately one log (746). The frequencies of wt-neutralizing and Beta-neutralizing sera were 58% and 12%, respectively (**Fig. 8c**). After a booster dose, 80% of sera from healthy donors were classified as Beta-neutralizing, and 31% as Omicron-neutralizing (**Fig. 8d**). The enhancement in responses after dose 3 was underestimated since the assay did not contain RBD from Omicron.

**Fig. 8.**
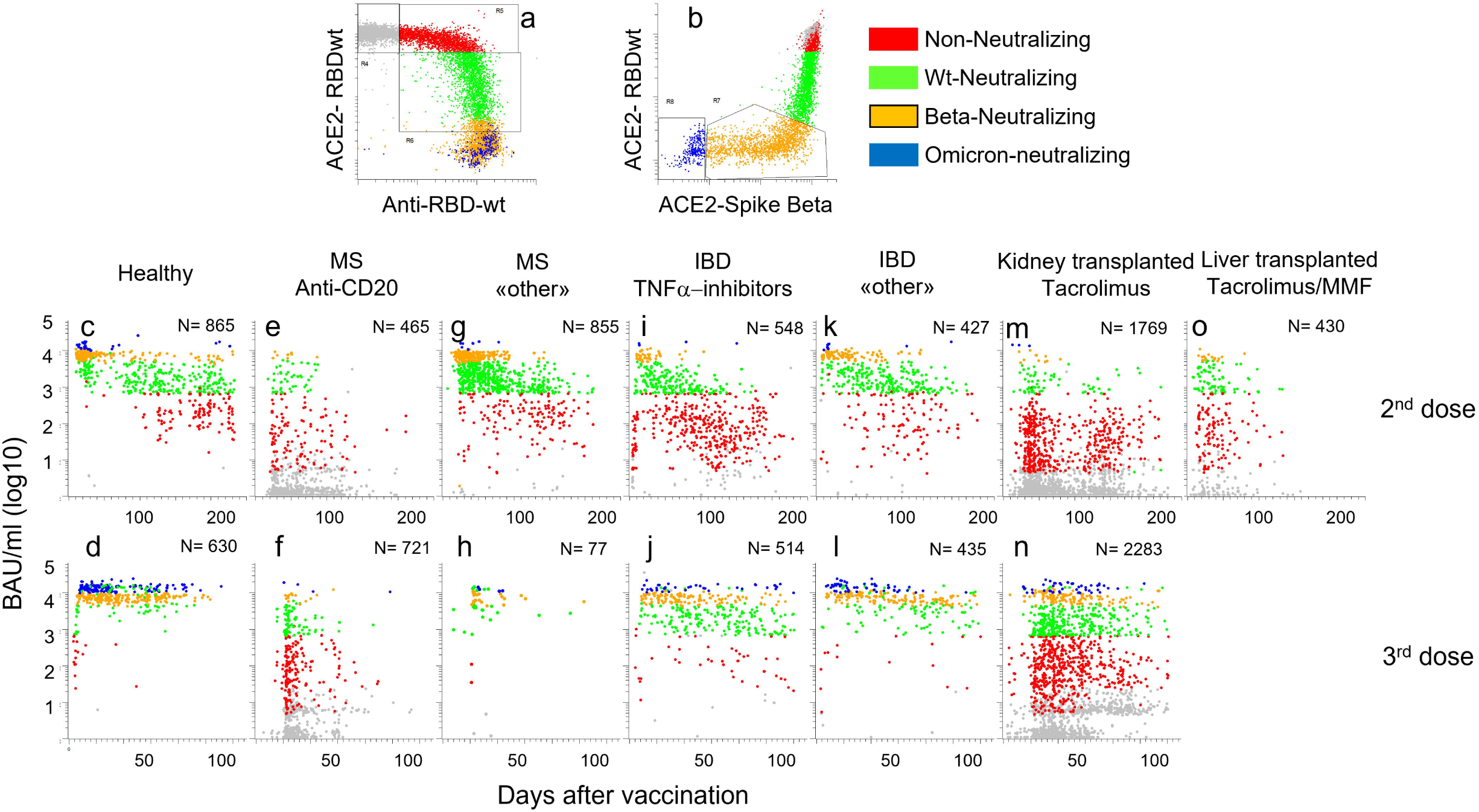
Large-scale analysis of humoral responses to COVID-19 vaccination. Indicated numbers of post-vaccine sera obtained from healthy individuals and patients on immunosuppressive therapy were analyzed with Multi-IgG-ACE2-RBD. Each dot corresponds to a different sample. **a-b:** see legend to Fig.3a-b. **c-n:** the x- and y-axes show days after vaccination and antibody levels in BAU/ml (log 10), respectively. MS: multiple sclerosis, IBD: Inflammatory bowel disease. “Other”: IBD: alpha 4 beta 7 antagonist, IL12_IL23 antagonist, MS: Cladribine, Leflunomide, glatirameracetate, interferon beta, Alpha 4 beta 1 antagonist. Tacrolimus was used in combination with other agents, mainly mycophenolic acid.

After two doses, 29% of 465 patients treated with anti-CD20 antibodies for multiple sclerosis (MS) had detectable antibodies, and 12% of sera were classified as “neutralizing” (**Fig. 8e**). Seroconversion increased to 44% after the 3^rd^ dose, but there was only a modest increase in the frequency of sera classified as wt-neutralizing (**Fig. 8e-f**). MS patients treated with Natalizumab, Cladribine, Glatirameracetate, or Leflunomide (**Fig. 8g-h, “other”**) had responses that were comparable to those observed in healthy individuals.

Treatment of inflammatory bowel disease with TNF-alpha antagonists (n=548) was associated with a shorter duration of the vaccine response (**Fig. 8i-j**). Thus, 80% of sera obtained earlier than 50 days after the 2^nd^ vaccine were classified as neutralizing, whereas the frequency fell to 15% after four months (**Fig. 8i-j**). At this time 13% had no detectable antibodies. This is in agreement with results from earlier studies ^32^. By comparison, patients who were treated for inflammatory bowel disease with antagonists for IL12/IL23 or α4β7 integrin (**Fig. 8k-l, “other”**) had responses that were comparable to those observed in healthy individuals.

Only 26 % of 1769 kidney transplant recipients treated with Tacrolimus had detectable antibodies to RBD-wt after two vaccines, and 6.5% of sera were classified as “wt-neutralizing” (**Fig. 8m**). However, a booster dose was quite effective in this group. More than half of the patients had detectable antibodies after dose 3, and nearly a third of sera were classified as neutralizing while 10% were Beta-neutralizing (**Fig. 8n**). The responses observed in sera from liver transplant recipients (n= 430) reflect the lower doses of Tacrolimus used in this patient cohort (**Fig. 8o**). Thus, 78% had detectable antibodies after two doses, and 40% were classified as “wt-neutralizing”. These results are in good agreement with those in earlier reports ^33^. Sera obtained after three doses were not available.

## DISCUSSION

We have used bead-based arrays to measure binding- and neutralizing antibodies to RBDs and Spike proteins from SARS-CoV-2wt and VOCs in 12,000 sera. The most important finding is that anti-RBD-wt titers correlate with the inhibitory effects of sera on ACE2-binding to RBDs from all VOCs tested (**Figs. 4-5**). The correlations between levels of binding antibodies could indicate that the most immunodominant epitopes are conserved in all variants (**Fig. 2**). RBD-ACE2 interaction assays, however, detect antibodies that bind to epitopes that undergo mutations driven by immune escape ^9-12^. Our results recapitulate those in earlier studies on differences in inhibitory effects of sera on ACE2-binding to RBDs and spike proteins from VOCs. An important new finding is that the effects followed a stringent pattern that was conserved across cohorts (**Fig. 3c**). Thus, there appears to be little inter-individual variation. Our cohort included healthy individuals as well as patients on a wide range of immunosuppressive therapies. We therefore conclude that anti-RBDwt titers have high predictive value for neutralizing activity against VOCs at the individual level.

Earlier studies have shown that neutralizing activity against the Omicron variant is almost exclusive to sera from individuals who have received a booster dose of COVID-19 vaccination virus ^19,34-37^. However, it has not been clear if the effect of a third vaccine dose on cross-protection is primarily qualitative or quantitative. Vaccination leads to time-dependent affinity maturation of B-cells and broadening of the epitope coverage ^8,20,38^. The ratio of serum antibodies that are capable of cross-neutralization may therefore increase after boosting ^20^. Here, we show that a quantitative difference in the antibody response is sufficient to explain the large increase in neutralization titers for Omicron observed after a booster dose. The results in **Fig. 6** predict that sera containing at least 11.000 BAU/ml have strong neutralizing effect against this VOC.

A simplistic interpretation of our results is that multiplexed assays are not needed since they yield essentially the same information as anti-RBD-wt titers. However, there are several good reasons to measure RBD-ACE2 interactions in parallel with anti-RBDwt titers. The two assays are orthogonal and yield independent evidence for the presence of RBD-binding antibodies. The methods are also complementary to the extent that they yield high resolution for high and low titers, respectively. A single dilution for each assay was sufficient to obtain a dynamic range of four logs. A high dynamic range usually comes at the cost of reduced throughput. For example, the ELISA protocol used to generate the results shown in **Fig. 5** involved duplicate measurement of each sample at eight serial dilutions. Multi-IgG-ACE2-RBD therefore yields a unique combination of high precision and throughput.

Multi-IgG-ACE2-RBD also opens the door to more standardized serology. Current diagnostic assays for anti-RBDwt titers are not interchangeable, even when results are converted to BAU/ml ^39^. Thus, the median titers reported for sera collected from healthy individuals after two doses of mRNA vaccines vary by more than seven-fold (959 to 7812 BAU/ml) ^1,2,6,7^. Variation in samples used as standards in different assays and laboratories is likely to be a contributing factor. Multi-IgG-ACE2-RBD eliminates the need for a standard. Instead, the relative binding of ACE2 to RBDs and spike proteins from VOCs serves as an internal reference. One may argue that membership in the groups identified in **Fig. 3a-b** only yields a rough estimate of anti-RBD titers. However, it is worth noting that the immune correlate for an absolute increase in vaccine efficacy of 10% was a 10-fold increase in titers of neutralizing antibodies ^1,2^ }. In this perspective, classification of sera on basis of membership in the groups identified here appears as a more robust alternative to numerical titers. The classification also seems more intuitive since there is a direct correlate to neutralizing activity.

The present study has limitations. First, we only measured serum neutralization of SARS-CoV-2wt. Further studies are needed to determine exact predictive values of Multi-IgG-RBD-ACE2 for protection against each VOC. Second, RBD-ACE2 interaction assays do not measure neutralizing antibodies binding outside of the RBD or factors that promote infectivity, such as virus replication rates. This may explain why the results shown in **Fig. 3c** did not recapitulate published differences in neutralizing titers for Delta and SARS-CoV-2wt (**Fig. 3c**) ^37,40^. The implication is that neutralization assays are still needed to establish if titers for new variants correlate with anti-RBD-wt levels ^3^. Finally, the arrays used for large-scale analysis of early vaccine responses shown in **Fig. 8**, did not contain RBD from Omicron. Reanalysis is necessary to determine if Omicron-neutralizing antibodies appear early, or first after many months of affinity maturation as suggested in a recent review ^20^. The results may have implications for establishing optimal dose intervals.

In conclusion, we show that anti-RBDwt titers in post-vaccine sera are broadly predictive for levels of neutralizing antibodies to VOCs. Our results also demonstrate the feasibility and utility of Multi-IgG-ACE2-RBD in large-scale immunomonitoring.

## Acknowledgements

This study was funded by grants from the Coalition of Epidemic Preparedness and Innovation (CEPI) and from South-Eastern Norway Regional Health Authority. The authors thank Prof. George Georgiou, University of Texas at Austin, USA for helpful discussions and critical reading of the manuscript.

## LEGENDS TO SUPPLEMENTARY FIGURES

**Fig S1:**
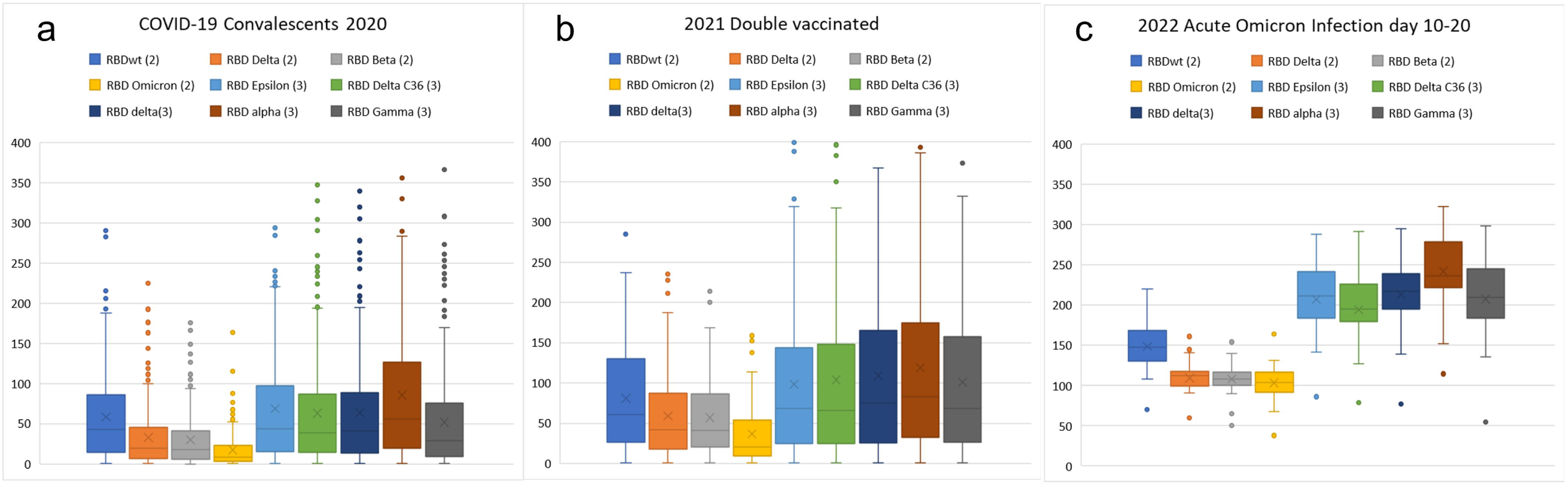
Signal values measured for antibodies to RBDs from SARS-CoV-2 variants. The bar blots show rMFI of anti-human IgG measured for beads coupled with RBD from indicated SARS-CoV-2 variant. Each dot corresponds to a different sample. Beads were incubated with serum diluted 1:1000 for 1h and labelled with anti-human IgG. Covid-19 convalescents: sera obtained in 2020 from individuals with mild COVID-19 (n=291). 2021 Double vaccinated: sera obtained in 2021 from individuals who had received two doses of the Pfizer/BioNTech vaccine (n=140). 2022 Acute Omicron Infection: sera obtained from healthy double-vaccinated individuals 10-20 days after symptom debut of Omicron infection (n=22).

**Fig. S2.**
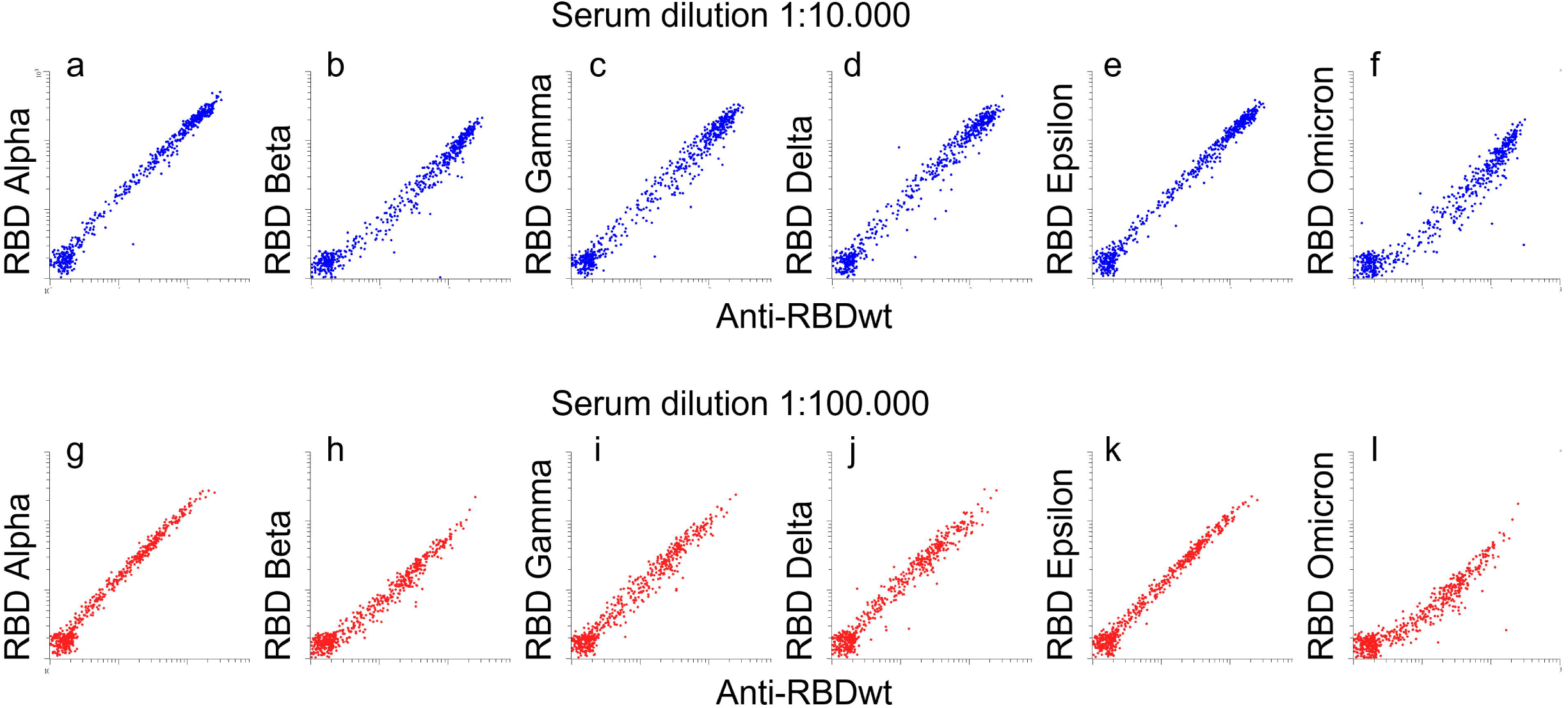
Correlation between antibody levels to RBDs from SARS-CoV-2 variants The dot plots show rMFI for anti-human IgG measured for beads coupled with RBDs from indicated SARS-CoV-2 variant. Each dot corresponds to a different sample. **a-f:** beads were incubated with 550 sera diluted 1:10,000 for 1h prior to labeling with anti-human IgG. **g-l:** beads were incubated with 550 sera diluted 1:100,000 for 1h prior to labeling with anti-human IgG.

**Fig. S3.**
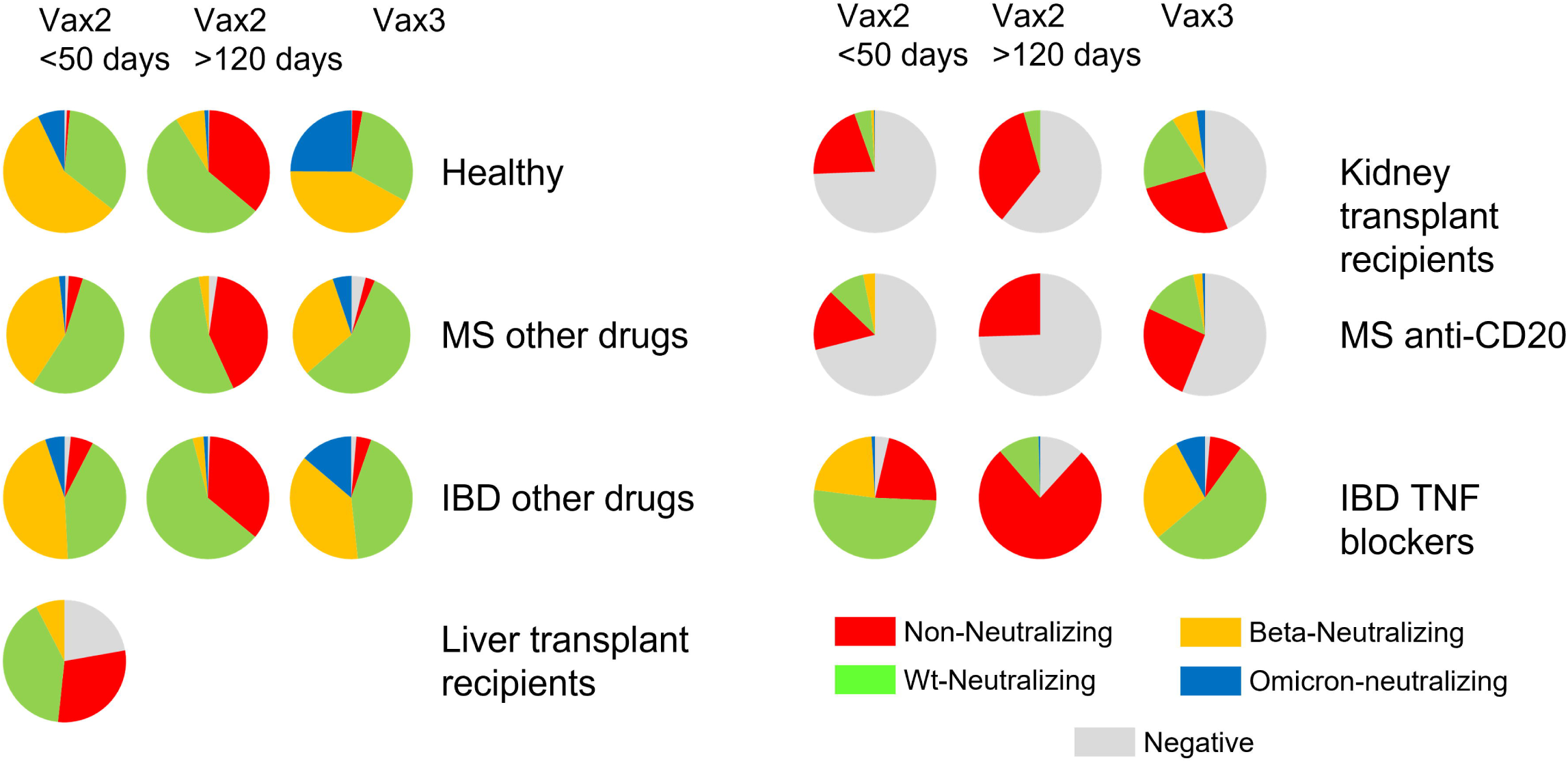
Classification of post-vaccine sera obtained from healthy individuals and patients on immunosuppressive therapy. The pie charts show frequencies of post-vaccine sera from indicated cohort classified according to group I-IV in Fig. 3a. See legend to Fig. 8.

